# Brain transcriptome analysis reveals gene expression differences associated with dispersal behaviour between range-front and range-core populations of invasive cane toads in Australia

**DOI:** 10.1101/2021.09.27.462079

**Authors:** Boris Yagound, Andrea J. West, Mark F. Richardson, Daniel Selechnik, Richard Shine, Lee A. Rollins

## Abstract

Understanding the mechanisms underlying rapid adaptation of invasive species in novel environments is key to improving our ability to manage these species. Many invaders demonstrate rapid evolution of behavioural traits involved in range expansion such as locomotor activity, exploration and risk-taking. However, the molecular mechanisms that underpin these changes are poorly understood. In 86 years, invasive cane toads (*Rhinella marina*) in Australia have drastically expanded their geographic range westward from coastal Queensland to Western Australia. During their range expansion, toads have undergone extensive phenotypic changes, particularly in behaviours that enhance the toads’ dispersal ability. Common-garden experiments have shown that some changes in behavioural traits related to dispersal are heritable. However, genetic diversity is greatly reduced across the invasive range due to a strong founder effect, and the genetic basis underlying dispersal-related behavioural changes remains unknown. Here we used RNA-seq to compare the brain transcriptomes of toads from the Hawai’ian source population, as well as three distinct populations from across the Australian invasive range. We found markedly different gene expression profiles between the source population and Australian toads. By contrast, cane toads from across the Australian invasive range had very similar transcriptomic profiles. Yet, key genes with functions putatively related to dispersal behaviour showed differential expression between range-core and range-front populations. These genes could play an important role in the behavioural changes characteristic of range expansion in Australian cane toads.

## Introduction

Invasive species are known for their high fecundity, phenotypic plasticity and excellent dispersal ability, which are all traits that enable them to adapt to and expand across novel environments (Prentis et al., 2008). As a result, invaders can pose significant threats to native species, ecosystem services, public health, agriculture and global economies (Clavero & Garcia-Berthou, 2005; Paini et al., 2016; Pimentel et al., 2005; Walsh et al., 2016). Behavioural traits are key to promoting range expansion of invasive species (Duckworth & Badyaev, 2007; Myles-Gonzalez et al., 2015; Rehage & Sih, 2004). Across taxa, dispersal success is associated with a suite of correlated behaviours. Individuals that are more active and more prone to taking risks (e.g., exploring unfamiliar habitats) have a greater probability of dispersing further and for longer periods of time (Duckworth & Badyaev, 2007; Myles-Gonzalez et al., 2015; Réale et al., 2007). Additionally, individuals with a greater propensity or aptitude for dispersal are more likely to be located on the expanding edge of populations than at the range-core, further increasing the overall dispersal rate (Duckworth & Badyaev, 2007; Phillips et al., 2006; Shine et al., 2011; Sol et al., 2002). The rate and extent of range expansion is influenced by the interplay between genetic factors, environmental factors, and phenotypic plasticity (Holway & Suarez, 1999).

Phenotypic variation between individuals is the substrate for selection on heritable traits, and is thus essential for evolution. Phenotypic variation in a population can result from genetic diversity or phenotypic plasticity. Because invasive populations often have low genetic diversity as a direct result of founder effects during introduction (Rollins et al., 2013), phenotypic plasticity may be especially important in these populations. Plasticity can affect behaviour, physiology and morphology (Piersma & Drent, 2003), and can evoke trait variation that could accelerate evolution (Robinson & Dukas, 1999). For example, increased plasticity at the leading edge of an expanding population could increase the expression of advantageous traits (e.g., locomotor ability and exploratory behaviour), which in turn can enhance the rate of dispersal. Phenotypic plasticity may thus be a key driver behind the rapid rates of range expansion common during invasions (Chuang & Peterson, 2016; Sexton et al., 2009).

The cane toad (*Rhinella marina*), native to South America, is a notorious invasive species now found across many parts of the world (Lever, 2001). In Australia, cane toads were introduced from a source population in Hawai’i to multiple sites along 1,200 km of Queensland coastline in 1935, in a failed attempt to control sugar cane beetles (Easteal, 1981; Shine, 2014; Shine et al., 2020). Since its introduction, the cane toad has expanded southward to New South Wales and westward through the Northern Territory, reaching Western Australia by 2010 (Shine, 2010; Urban et al., 2008). Across the western half of that invasion trajectory, climatic conditions are hotter and seasonally much more arid compared to both the toad’s native range and its Queensland introduction sites (Kearney et al., 2008; Kosmala et al., 2020). Moreover, substantial variation in morphology, physiology and behaviour linked to dispersal ability have been documented across the toads’ westward invasion in Australia (Shine, 2010). For example, toads located on the invasion front have longer legs (Phillips et al., 2006), wider forelimbs, narrower hindlimbs, more compact skulls (Hudson et al., 2016), smaller relative head widths (Hudson et al., 2018), greater endurance (Llewelyn et al., 2010), move more frequently (Alford et al., 2009) and further in a given period of time (Lindstrom et al., 2013), and are more exploratory and more likely to exhibit risk-taking behaviour in a novel environment than are individuals from range-core populations (Gruber et al., 2017). These phenotypic changes have culminated in increased dispersal ability in toads from range-front populations (Urban et al., 2008).

Despite the breadth of phenotypic variation in invasive Australian cane toads, genetic variation is low (Lillie et al., 2017; Selechnik et al., 2019a; Slade & Moritz, 1998). Interestingly, although genome-wide genetic diversity in Australia is overall greatly reduced as compared to the native range, diversity is increased at some loci that appear to be under selection and that are associated with climatic conditions (Selechnik et al., 2019a). Additionally, common-garden experiments have shown that some of the phenotypic traits showing extensive variation across Australia are heritable (e.g. morphological, (Hudson et al., 2016; Hudson et al., 2018); physiological, (Brown et al., 2015; Kosmala et al., 2018); or behavioural, (Gruber et al., 2017; Stuart et al., 2019)). Further, muscle and spleen transcriptome analyses found hundreds of differentially expressed genes (hereafter, DEGs) between range-core and range-front populations. These genes are putatively involved in metabolism, cellular repair and immune function, which might indicate a molecular response to greater environmental stressors at the range edge (Rollins et al., 2015; Selechnik et al., 2019b). Overall, this collection of evidence indicates a genetic basis underlying the significant phenotypic variation seen in cane toads across the Australian invasive range.

With respect to plasticity in Australian cane toads, studies have yielded mixed results. Less plasticity in growth and development was found in individuals from the western range-front *versus* those from the range-core (Ducatez et al., 2016). When juvenile toads were exposed to high *versus* low exercise regimes, body size in range-front toads was less plastic compared to that of range-core toads, but this trend was reversed in toads whose diets were supplemented with calcium (Stuart et al., 2019). Greater plasticity with respect to temperature tolerance was found in toads from the southern range-front *versus* those from the range-core (Kolbe et al., 2010; McCann et al., 2014). These studies demonstrate that patterns of plasticity in morphological and physiological traits are complex in toads sampled across the Australian invasive range. To date, little is known about behavioural plasticity in these populations.

Here we used RNA-seq to analyse brain transcriptomes of toads sampled from Hawai’i, the area from which toads were taken to Australia, as well as three geographically distinct populations from across the Australian invasive range. This sampling design allowed us to identify candidate genes underlying range expansion traits in Australian cane toads and to explore differences in variation in gene expression (i.e., plasticity) across these sampling sites.

## Materials and Methods

### Sample collection

We collected brain tissue from a total of 54 wild, adult female cane toads during April and May of 2014 and 2015 from 13 sites (N = 4/5 individuals per site) representing four populations in Hawai’i and Australia (Figure 1 and Table S1). Populations corresponded to distinct genetic clusters across the invasive range following Selechnik et al. (2019): source (Hawai’i), range-core (coastal Queensland), intermediate (central Queensland and Northern Territory) and range-front (Western Australia) (Figure 1). Immediately following capture, we humanely euthanized toads, extracted their brains and stored them in RNAlater (Qiagen, USA) at 4°C to preserve tissue integrity until samples were transported to the lab, where they were stored at −80°C prior to RNA extraction. All experimental procedures involving live toads were approved by the University of Sydney Animal Care and Ethics Committee (2014/562) and the Deakin University Animal Ethics Committee (AEX04-2014).

**Figure 1.**
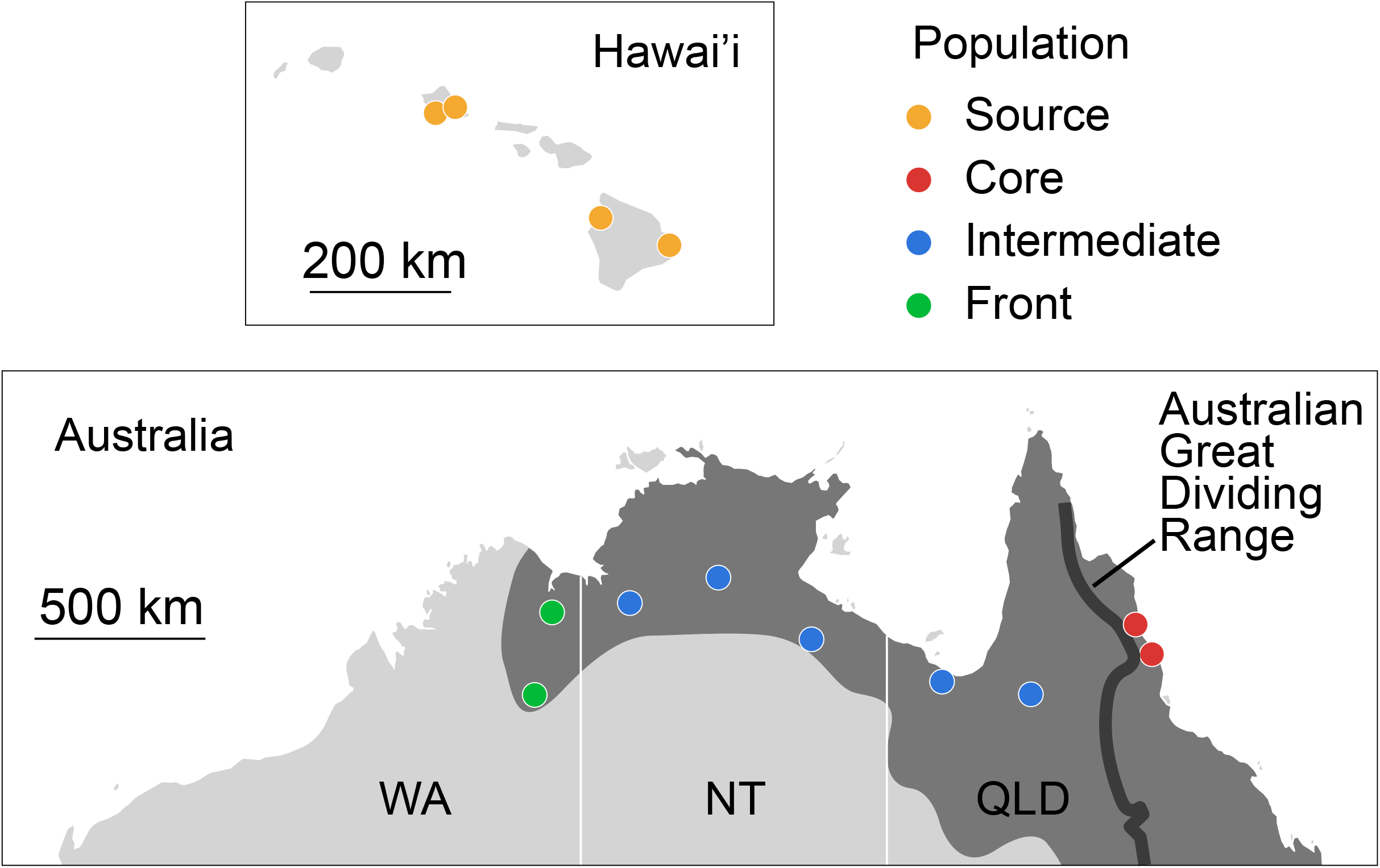
Location of samples. QLD, Queensland; NT, Northern Territory; WA, Western Australia. The shaded area represents the cane toad’s Australian invasive range.

### RNA extraction and sequencing

We extracted total RNA using Qiagen RNeasy Lipid Tissue Mini Kits (Qiagen), following the manufacturer’s protocol. We homogenised tissues using a Fast Prep-24 Classic homogeniser (MP Biomedicals, USA) and 1 mm Zirconia/Silica beads (Daintree Scientific, Australia) for 1 minute at 6 m/s. We digested genomic DNA using a Qiagen RNase-Free DNase set on column during extraction. We quantified extracted RNA using a Qubit RNA HS assay kit on a Qubit 3.0 Fluorometer (Life Technologies, USA). Library preparation and sequencing was conducted commercially at Macrogen (South Korea). Library preparation followed the TruSeq mRNA 2 (Illumina, USA) protocol and libraries were sequenced on an Illumina HiSeq 2500 platform (two lanes of 125 bp paired-end sequencing), generating 872 million reads.

### Data pre-processing, alignment and gene expression quantification

We used FastQC 0.11.7 (http://www.bioinformatics.babraham.ac.uk/projects/fastqc) to check the quality of the raw data. Trimmomatic 0.38 (Bolger et al., 2014) was used to remove adaptor sequences and trim low quality reads with the following parameters: ILLUMINACLIP: path/to/TruSeq3-PE.fa:2:30:10:4 HEADCROP:13 AVGQUAL:30 MINLEN:36. We mapped trimmed reads to the multi-tissue reference cane toad transcriptome (Richardson et al., 2018) using STAR 2.7.2b (Dobin et al., 2013) in two-pass mode with default parameters. We used resultant BAM files to quantify gene expression using Salmon 1.2.1 (Patro et al., 2017).

### Differential expression analysis

We used edgeR 3.32.1 (Robinson et al., 2010) to filter out genes using the filterByExpr function with default parameters in R 4.0.4 (R Core Team, 2021). We further filtered out genes that had less than 10 counts per million in at least 10 samples. To assess the presence of outliers, we normalised and rlog-transformed counts before computing pairwise correlation for all the samples. We then used the resultant correlation matrix to plot a heatmap with pheatmap 1.0.12 (Kolde, 2019). This revealed two outliers (Figure S1), B24 and B31, that we excluded from subsequent analyses. We performed differential expression analysis using DESeq2 1.30.1 (Love et al., 2014). We performed differential expression analysis between the source population and each population of the Australian range, and second between all three populations of the Australian range. We considered all genes with Benjamini-Hochberg adjusted *p*-values < 0.05 (Benjamini & Hochberg, 1995) to be significantly differentially expressed between any pairwise comparison. We further performed differential expression analysis using a likelihood ratio test with DESeq2 between all nine Australian sites ordered along an east-to-west transect. This transect reflects the timeline of the cane toad invasion and was aimed at testing whether differences in brain gene expression followed a continuum across the Australian range rather than population-specific changes. We then used the degPatterns function in DESeq2 on all DEGs across this transect to identify clusters of genes with similar expression profiles across the Australian range. This tool performs a hierarchical clustering based on gene pairwise Kendall correlations.

### Network analysis

We used WGCNA 1.70-3 (Langfelder & Horvath, 2008) to perform a gene correlation network analysis and test whether distinct clusters of coregulated genes can be identified along the Australian invasive range. We used the pickSoftThreshold function to select a soft threshold power according to the authors recommendations (Zhang & Horvath, 2005). We used the blockwiseModules function with power = 10 to identify gene coexpression modules. We then fitted linear models to test whether each module was significantly associated with any of the 3 populations of the Australian range, using limma 3.46.0 (Ritchie et al., 2015) with Benjamini-Hochberg correction for multiple testing (same as above).

### Differential variability analysis

We used MDSeq 1.0.5 (Ran & Daye, 2017) to test whether brain genes differed in their expression variability (hereafter, dispersion) across the Australian range. Gene expression variability is being recognised as an important driver of phenotypic differences (Ecker et al., 2018). We conducted differential dispersion analysis between the three Australian populations (i.e. range-core, intermediate and range-front). We randomly selected two sites (Croydon and Mataranka) of the intermediate population for this analysis to balance sample sizes with the range-core and range-front populations. We normalised gene counts using the trimmed mean of M-values (TMM) method (Robinson & Oshlack, 2010) in edgeR. Any gene with a Benjamini-Hochberg adjusted *p*-value < 0.05 was considered to be significantly differentially dispersed between any pairwise comparison.

### Functional analysis

We performed Gene Ontology (GO) analysis using goseq 1.26.0 (Young et al., 2010) to infer biological function of all differentially expressed or differentially dispersed gene sets. We conducted GO analysis using the probability weighting function (PWF) to adjust for transcript length bias and the Wallenius approximation to test for over-representation. *P*-values were adjusted with the Benjamini-Hochberg method. We visualised GO results using enrichplot (Yu, 2021).

## Results

Cane toads from the Australian range showed extensive differences in brain gene expression compared to toads from the Hawaiian source population. There were 6,529 DEGs between range-core and source populations, 7,769 DEGs between intermediate and source populations, and 6,770 DEGs between range-front and source populations (Figure 2A–F). The number of upregulated genes was higher in each Australian population compared to the source population (Wilcoxon test: *p* = 0.040). A core set of 4,903 genes were commonly differentially expressed between the source population and each Australian population (Figure 2G). GO analysis of these 4,903 DEGs revealed significant enrichment for processes such as translation and mitochondrial function (Figure 2H).

**Figure 2.**
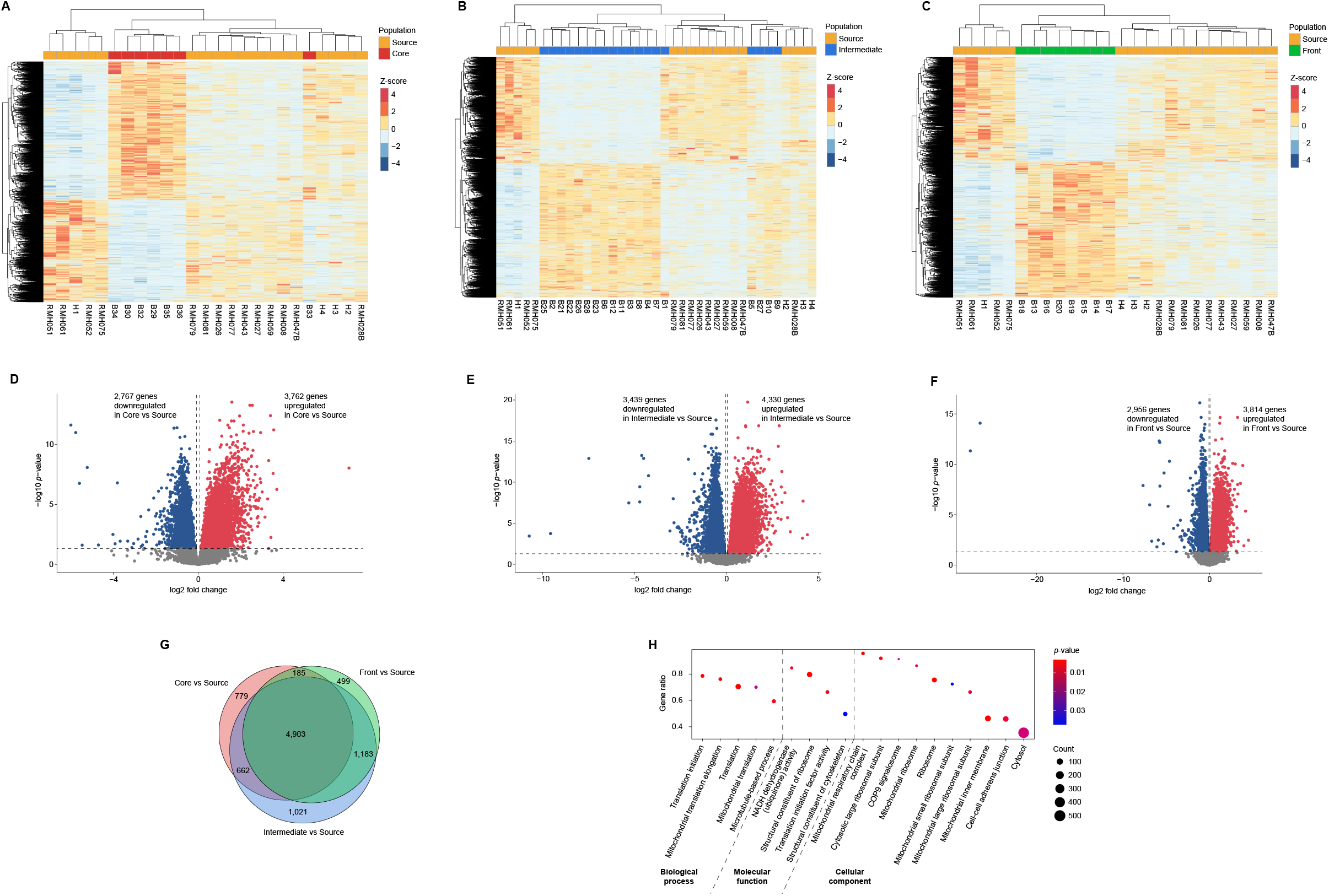
(**A–C**) Heatmap of normalised gene expression values for all DEGs between (**A**) core and source populations, (**B**) intermediate and source populations, and (**C**) front and source populations. Columns correspond to samples. Rows correspond to genes. Colour depicts Z-score normalised gene expression value. (**D–F**) Volcano plots of significantly DEGs between (**D**) core and source populations, (**E**) intermediate and source populations, and (**F**) front and source populations. Nonsignificant genes are represented in grey. (**G**) Overlap of DEGs across each pairwise comparison. (**H**) GO analysis of the core set of DEGs overlapping across each pairwise comparison. The size of each circle is proportional to the number of genes being significantly enriched, while the colour of each circle is proportional to its FDR-corrected *p*-value. Gene ratio corresponds to the proportion of genes being enriched out of the total number of genes in that GO category.

By contrast, cane toads within the Australian range showed few geographic differences in brain gene expression. There were only 59 DEGs between intermediate and range-core populations (Table 1 and Figure 3A,C), and 21 DEGs between range-front and range-core populations (Table 2 and Figure 3B,D). Nine genes were commonly differentially expressed in intermediate versus range-core populations and range-front versus range-core populations (Figure 3E). Specifically, *F12*, *FANCD2* and *POL* were upregulated in intermediate and range-front populations compared to range-core populations, while *POMC*, *L1RE1*, *MAN2B1* and *POL4* were downregulated in intermediate and range-front populations compared to range-core populations. There was no significant enrichment for GO terms in any pairwise comparison. Range-front and intermediate populations showed no significant differences in brain gene expression.

**Figure 3.**
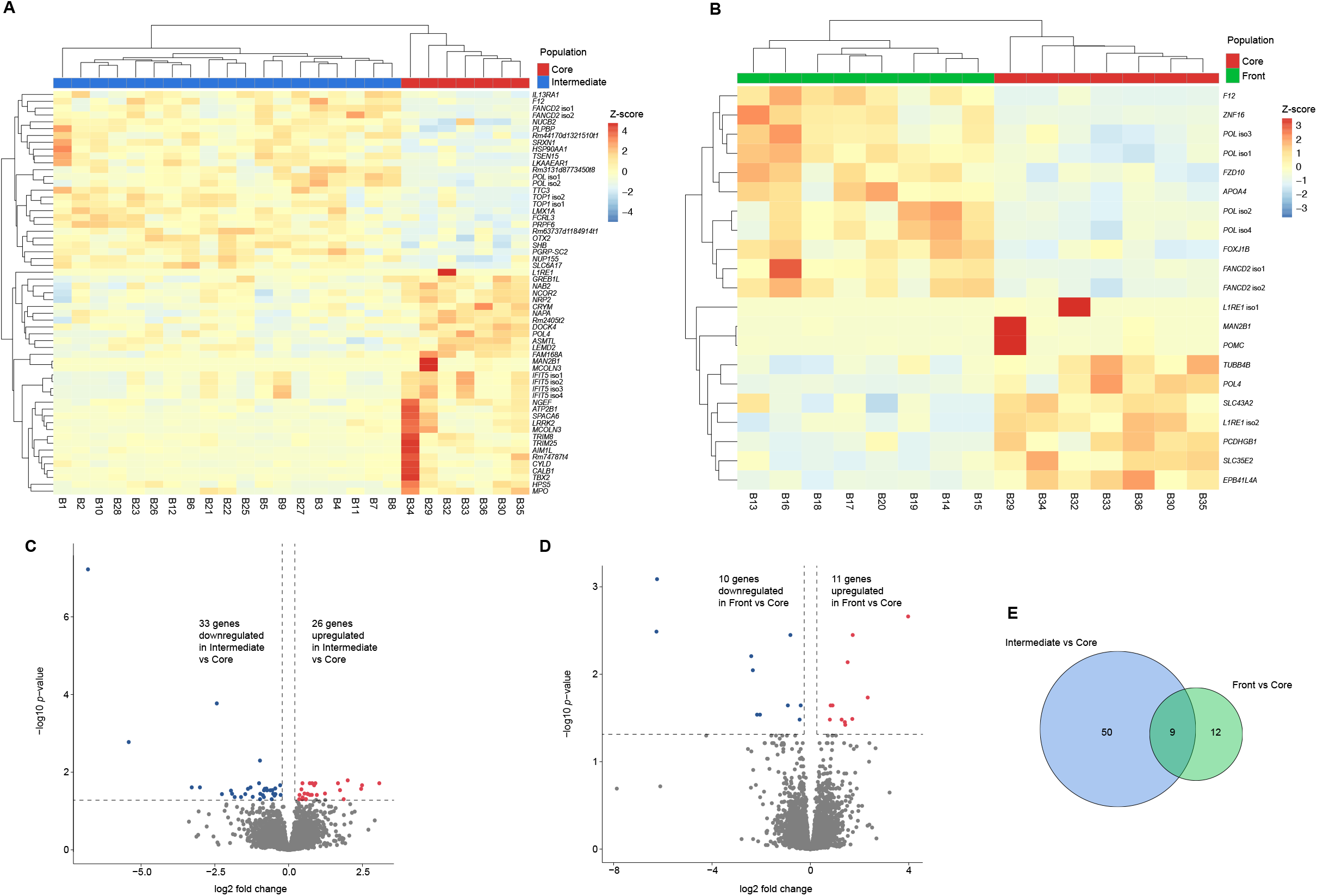
(**A,B**) Heatmap of normalised gene expression values for all DEGs between (**A**) intermediate and core populations, and (**B**) front and core populations. Columns correspond to samples. Rows correspond to genes. Colour depicts Z-score normalised gene expression value. (**C,D**) Volcano plots of significantly DEGs between (**C**) intermediate and core populations, and (**D**) front and core populations. Nonsignificant genes are represented in grey. (**E**) Overlap of DEGs across each pairwise comparison.

**Table 1.** DEGs between intermediate and core populations. Genes respectively upregulated and downregulated in intermediate vs core populations are indicated by log2 fold change values respectively > 0 and < 0.

| Gene | Protein | Log2 fold change | p-value |
| --- | --- | --- | --- |
| *POMC* | Pro-opiomelanocortin | -6.80 | < 0.00001 |
| *MAN2B1* | Lysosomal alpha-mannosidase | -2.43 | 0.0002 |
| *L1RE1* | LINE-1 retrotransposable element ORF2 protein | -5.42 | 0.0018 |
| *CRYM* | Ketimine reductase mu-crystallin | -0.97 | 0.0050 |
| *TTC3* | E3 ubiquitin-protein ligase TTC3 isoform X4 | 2.00 | 0.0163 |
| *F12* | Coagulation factor XII | 3.07 | 0.0192 |
| *TOP1* (isoform 1) | DNA topoisomerase 1 | 1.67 | 0.0193 |
| *TOP1* (isoform 2) | DNA topoisomerase 1 | 0.91 | 0.0193 |
| *FANCD2* (isoform 1) | Fanconi anemia group D2 protein | 0.80 | 0.0193 |
| *POL* (isoform 1) | Pol polyprotein | 0.72 | 0.0193 |
| *NUP155* | Nuclear pore complex protein Nup155 | 0.47 | 0.0193 |
| *FAM168A* | Protein FAM168A | -1.00 | 0.0193 |
| *LKAAEAR1* | Protein LKAAEAR1 | 2.49 | 0.0217 |
| *NUCB2* | Nucleobindin-2 | 0.87 | 0.0217 |
| *NCOR2* | Nuclear receptor corepressor 2 | -0.29 | 0.0217 |
| *TRIM25* | E3 ubiquitin/ISG15 ligase TRIM25 | -1.29 | 0.0247 |
| *CALB1* | Calbindin | -3.00 | 0.0247 |
| *Rm74787t4* | Unknown | -3.28 | 0.0247 |
| *NRP2* | Neuropilin-2 | -0.46 | 0.0263 |
| *IFIT5* (isoform 1) | Interferon-induced protein with tetratricopeptide repeats 5 | -0.75 | 0.0265 |
| *Rm2405t2* | Unknown | -0.81 | 0.0268 |
| *Rm3131d8773450t8* | Unknown | 2.46 | 0.0269 |
| *SPACA6* | Sperm acrosome membrane-associated protein 6 | -1.38 | 0.0269 |
| *PLPBP* | Proline synthase co-transcribed bacterial homolog protein | 0.43 | 0.0277 |
| *Rm63737d1184914t1* | Unknown | 1.74 | 0.0291 |
| *LEMD2* | LEM domain-containing protein 2 | -0.55 | 0.0291 |
| *HPS5* | Hermansky-Pudlak syndrome 5 protein homolog | -0.62 | 0.0293 |
| *IFIT5* (isoform 2) | Interferon-induced protein with tetratricopeptide repeats 5 | -0.73 | 0.0293 |
| *IFIT5* (isoform 3) | Interferon-induced protein with tetratricopeptide repeats 5 | -0.83 | 0.0293 |
| *MCOLN3* | Mucolipin-3 | -1.96 | 0.0299 |
| *FANCD2* (isoform 2) | Fanconi anemia group D2 protein | 1.23 | 0.0355 |
| *SLC6A17* | Sodium-dependent neutral amino acid transporter SLC6A17 | 0.54 | 0.0355 |
| *FCRL3* | Fc receptor-like protein 3 | 0.64 | 0.0361 |
| *ASMTL* | N-acetylserotonin O-methyltransferase-like protein | −0.52 | 0.0361 |
| *GREB1L* | GREB1-like protein | −0.96 | 0.0361 |
| *MPO* | Myeloperoxidase | −1.92 | 0.0361 |
| *POL* (isoform 2) | Pol polyprotein | 0.65 | 0.0367 |
| *DOCK4* | Dedicator of cytokinesis protein 4 | −0.45 | 0.0367 |
| *TBX2* | T-box transcription factor TBX2 | −1.47 | 0.0367 |
| *AIM1L* | Absent in melanoma 1-like protein | −2.26 | 0.0367 |
| *Rm44170d1321510t1* | Unknown | 0.79 | 0.0385 |
| *OTX2* | Orthodenticle homeobox protein OTX2 | 0.96 | 0.0386 |
| *LMX1A* | LIM homeobox transcription factor 1-alpha | 0.72 | 0.0386 |
| *PRPF6* | Pre-mRNA-processing factor 6 | 0.37 | 0.0386 |
| *NAPA* | Alpha-soluble NSF attachment protein | −0.27 | 0.0386 |
| *IFIT5* (isoform 4) | Interferon-induced protein with tetratricopeptide repeats 5 | −0.87 | 0.0392 |
| *NGEF* | Ephexin-1 | −0.48 | 0.0401 |
| *TRIM8* | Probable E3 ubiquitin-protein ligase TRIM8 | −1.22 | 0.0433 |
| *POL4* | Retrovirus-related Pol polyprotein from transposon 412 | −1.61 | 0.0438 |
| *ATP2B1* | Plasma membrane calcium-transporting ATPase 1 | −0.84 | 0.0438 |
| *LRRK2* | Leucine-rich repeat serine/threonine-protein kinase 2 | −1.83 | 0.0438 |
| *SHB* | SH2 domain-containing adapter protein B | 0.48 | 0.0449 |
| *PGRP-SC2* | Peptidoglycan-recognition protein SC2 | 1.87 | 0.0497 |
| *SRXN1* | Sulfiredoxin-1 | 0.59 | 0.0497 |
| *IL13RA1* | Interleukin-13 receptor subunit alpha-1 | 0.59 | 0.0497 |
| *TSEN15* | tRNA-splicing endonuclease subunit Sen15 | 0.46 | 0.0497 |
| *HSP90AA1* | Heat shock protein HSP 90-alpha | 0.36 | 0.0497 |
| *NAB2* | NGFI-A-binding protein 2 | −0.57 | 0.0497 |
| *CYLD* | Ubiquitin carboxyl-terminal hydrolase CYLD | −0.97 | 0.0497 |

**Table 2.** DEGs between front and core populations. Genes respectively upregulated and downregulated in front vs core populations are indicated by log2 fold change values respectively > 0 and < 0.

| Gene | Protein | Log2 fold change | p-value |
| --- | --- | --- | --- |
| *POMC* | Pro-opiomelanocortin | -6.24 | 0.0008 |
| *F12* | Coagulation factor XII | 3.98 | 0.0022 |
| *L1RE1* (isoform 1) | LINE-1 retrotransposable element ORF2 protein | -6.26 | 0.0033 |
| *POL* (isoform 1) | Pol polyprotein | 1.72 | 0.0036 |
| *L1RE1* (isoform 2) | LINE-1 retrotransposable element ORF2 protein | -0.81 | 0.0036 |
| *MAN2B1* | Lysosomal alpha-mannosidase | -2.40 | 0.0062 |
| *APOA4* | Apolipoprotein A-IV | 1.52 | 0.0073 |
| *SLC35E2* | Solute carrier family 35 member E2 | -2.34 | 0.0090 |
| *ZNF16* | Zinc finger protein 16 | 2.33 | 0.0185 |
| *FANCD2* (isoform 1) | Fanconi anemia group D2 protein | 0.90 | 0.0227 |
| *POL* (isoform 2) | Pol polyprotein | 0.82 | 0.0227 |
| *EPB41L4A* | Erythrocyte membrane protein band 4.1-like protein 4A | -0.92 | 0.0227 |
| *TUBB4B* | Tubulin beta-4B chain | -0.39 | 0.0227 |
| *POL4* | Retrovirus-related Pol polyprotein from transposon 412 | -2.17 | 0.0290 |
| *PCDHGB1* | Protocadherin gamma-B1 | -2.05 | 0.0290 |
| *POL* (isoform 3) | Pol polyprotein | 1.70 | 0.0325 |
| *FOXJ1B* | Forkhead box protein J1-B | 1.27 | 0.0330 |
| *POL* (isoform 4) | Pol polyprotein | 0.79 | 0.0330 |
| *SLC43A2* | Large neutral amino acids transporter small subunit 4 | -0.44 | 0.0330 |
| *FZD10* | Frizzled-10-A | 1.40 | 0.0353 |
| *FANCD2* (isoform 2) | Fanconi anemia group D2 protein | 1.42 | 0.0379 |

There were 228 DEGs across all nine Australian sites ordered along an east-to-west transect (top 50 DEGs are shown in Table 3). Of these, 31 genes had been previously identified in this study as showing significant differences in expression between populations, including *POMC*, *MAN2B1*, *CALB1*, *NUCB2* and *LRRK2*. Among these 228 DEGs, we identified four clusters of genes showing similar expression profiles across the Australian range. These four clusters showed curvilinear patterns of gene expression whereby gene expression in intermediate areas differed from that in range-core and range-front areas (Figure 4A-D). Clusters 2 and 3 showed significant enrichment for GO terms such as RNA-mediated transposition, cyclooxygenase pathway and oxidation-reduction process (Figure 4E,F).

**Figure 4.**
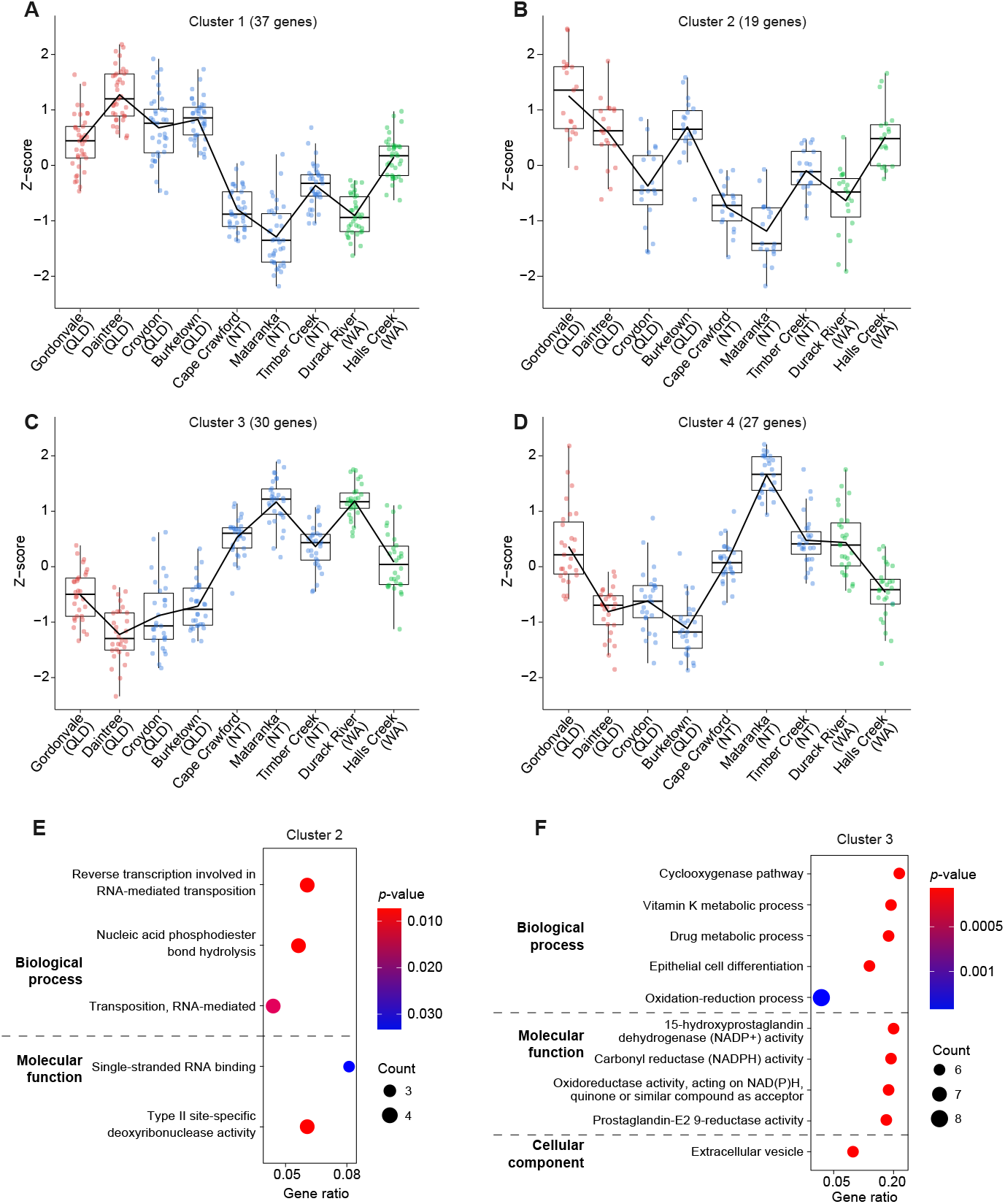
(**A–D**) Z-score abundance of gene expression of DEGs across the Australian range. Sites on the x-axis are ordered from east to west. Genes showing similar patterns of gene expression are grouped together. Box plots represent median, interquartile range and 95% confidence interval. Black lines represent the trend in expression change. Colours correspond to populations (red, core; blue, intermediate; green, front). (**E,F**) GO analysis of DEGs belonging to clusters 2 (**E**) and 3 (**F**). The size of each circle is proportional to the number of genes being significantly enriched, while the colour of each circle is proportional to its FDR-corrected *p*-value. Gene ratio corresponds to the proportion of genes being enriched out of the total number of genes in that GO category.

**Table 3.**
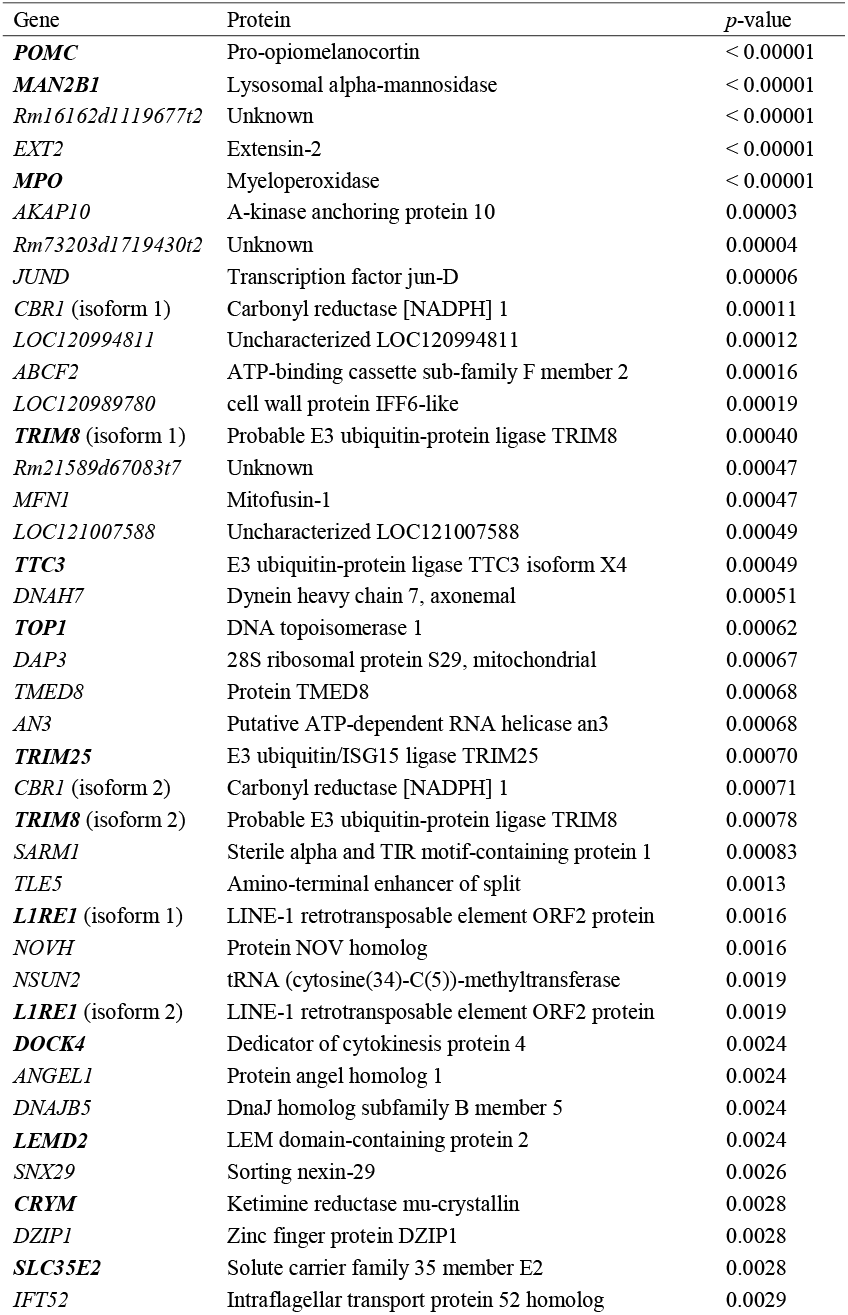
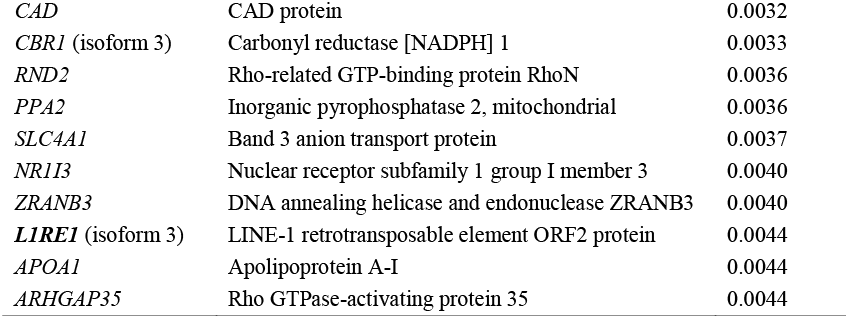
Top 50 genes showing significant expression changes across the whole Australian range. Genes in bold also show significant expression differences between populations.

Gene correlation network analysis revealed 68 modules of coregulated genes. Yet, no module was significantly associated with the toads’ location across the Australian range (linear models: all *p* > 0.133).

Toads within the Australian range also showed moderate geographic differences in the variability of brain gene expression. There were 17 differentially dispersed genes between intermediate and range-core populations (Table 4 and Figure 5A), 14 differentially dispersed genes between range-front and range-core populations (Table 5 and Figure 5B), and 29 differentially dispersed genes between range-front and intermediate populations (Table 6 and Figure 5C). The number of over-dispersed genes was similar in both populations in each pairwise comparison (Tables 4–6 and Figure 5A–C). Thus, changes in gene expression variability were not characteristic of any population within the Australian range. Five genes were commonly differentially dispersed in intermediate versus range-core populations and range-front versus range-core populations (Figure 5D), namely *WRB*, *MAN2B1*, *POL*, *PAK2* and *L1RE1*. Four genes were commonly differentially dispersed in range-front versus intermediate populations and intermediate versus range-core populations (Figure 5D), namely *MCOLN3*, *LRRK2* and two uncharacterised genes. One gene, *NPW*, was commonly differentially dispersed in range-front versus intermediate populations and range-front versus range-core populations (Figure 5D). No gene was commonly differentially dispersed in all three pairwise comparisons. There was no significant enrichment for any GO term in any pairwise comparison. Six genes, *MCOLN3*, *MAN2B1*, *LRRK2*, *EPB41L4A*, *NUCB2* and *L1RE1*, were both differentially expressed and differentially dispersed between at least two populations (Figure 5E).

**Figure 5.**
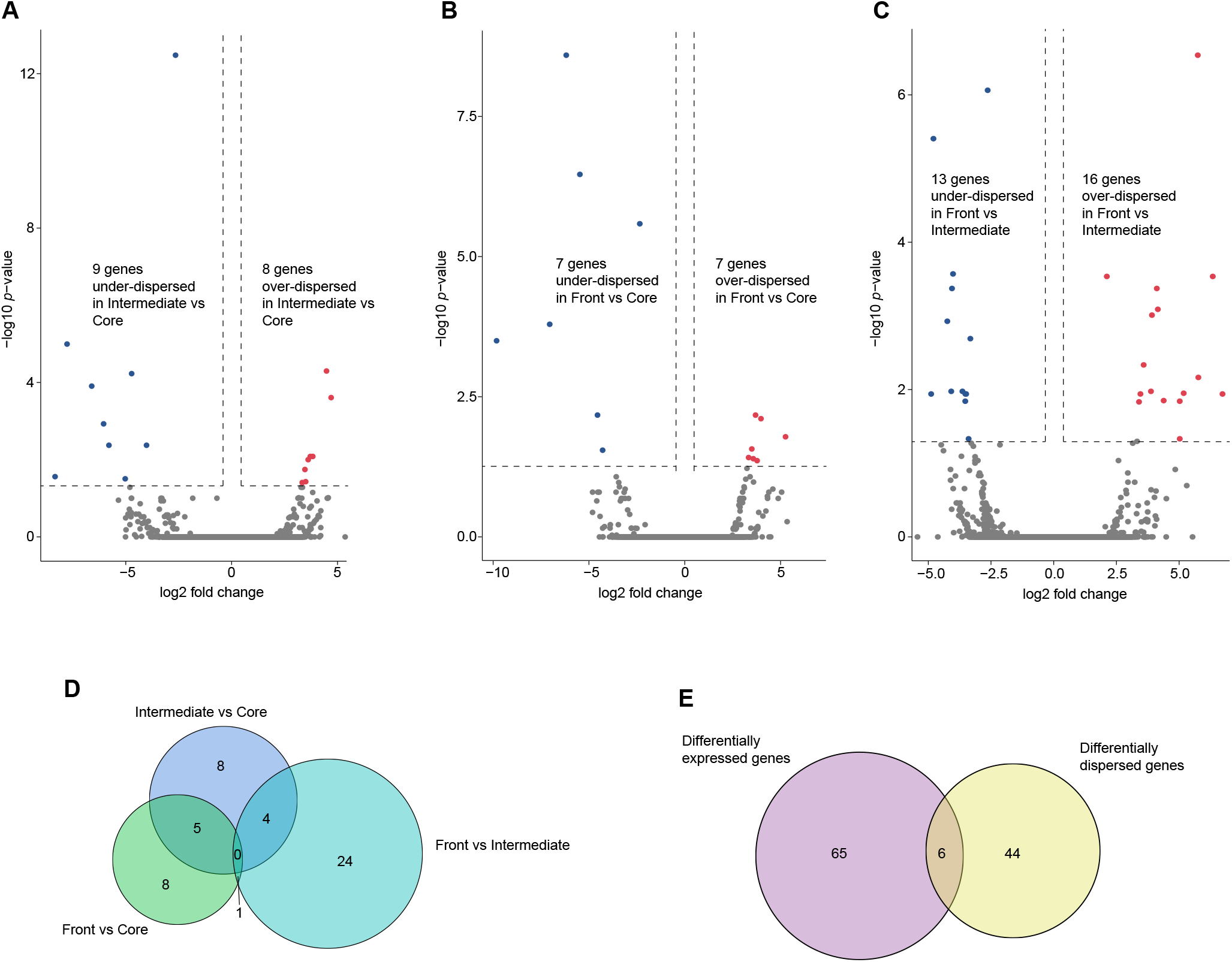
(**A–C**) Volcano plots of significantly differentially dispersed genes between (**A**) intermediate and core populations, (**B**) front and core populations, and (**C**) front and intermediate populations. Nonsignificant genes are represented in grey. (**D**) Overlap of differentially dispersed genes across each pairwise comparison. (**E**) Overlap of differentially expressed genes and differentially dispersed genes across all pairwise comparisons.

**Table 4.**
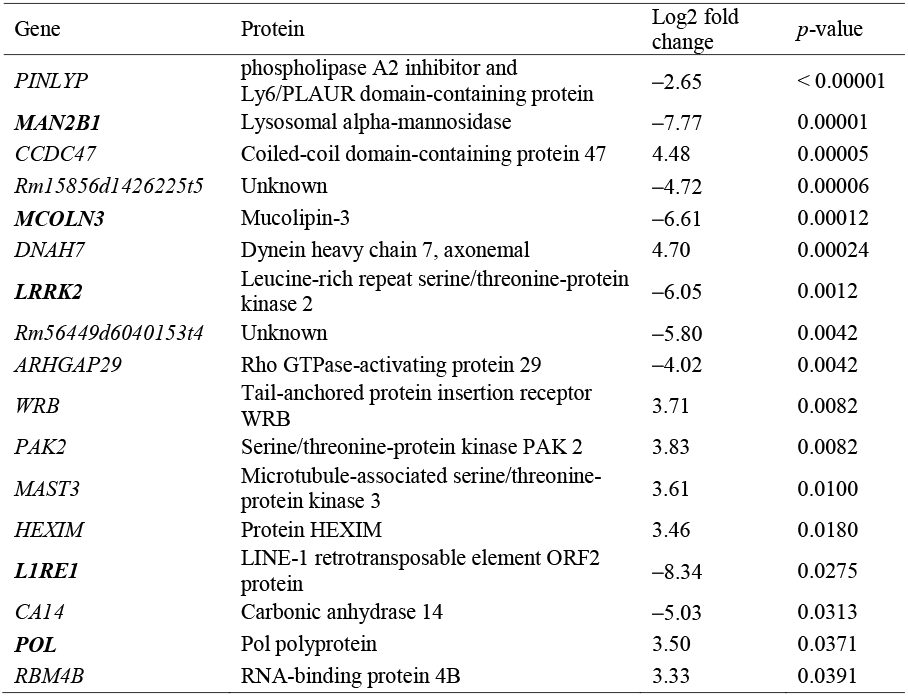
Differentially dispersed genes between intermediate and core populations. Genes respectively under-dispersed and over-dispersed in intermediate vs core populations are indicated by log2 fold change values respectively > 0 and < 0. Genes in bold also show significant expression differences between populations.

**Table 5.**
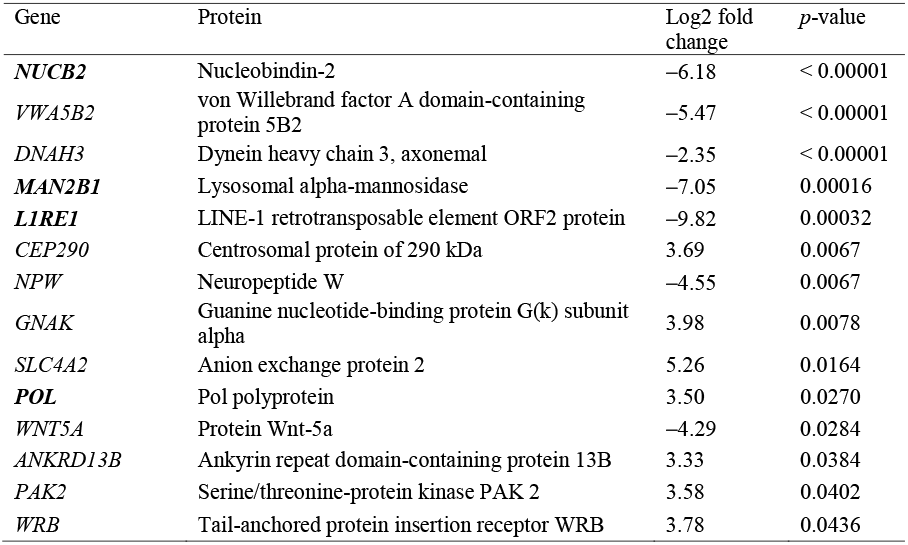
Differentially dispersed genes between front and core populations. Genes respectively under-dispersed and over-dispersed in front vs core populations are indicated by log2 fold change values respectively > 0 and < 0. Genes in bold also show significant expression differences between populations.

**Table 6.** Differentially dispersed genes between front and intermediate populations. Genes respectively under-dispersed and over-dispersed in front vs intermediate populations are indicated by log2 fold change values respectively > 0 and < 0. Genes in bold also show significant expression differences between populations.

| Gene | Protein | Log2 fold change | p-value |
| --- | --- | --- | --- |
| *AGRP* | Agouti-related protein | 5.74 | $< 0.00001$ |
| *SAMD10* | Sterile alpha motif domain-containing protein 10 | -2.63 | $< 0.00001$ |
| *KRAS* | GTPase KRas | -4.80 | $< 0.00001$ |
| *PPP2R5B* | Serine/threonine-protein phosphatase 2A 56 kDa regulatory subunit beta isoform | -4.01 | 0.00027 |
| *TNFAIP2* | Tumor necrosis factor alpha-induced protein 2 | 2.12 | 0.00029 |
| ***MCOLN3*** | Mucolipin-3 | 6.34 | 0.00029 |
| *SH2D4A* | SH2 domain-containing protein 4A | 4.11 | 0.00042 |
| *NUMA1* | Nuclear mitotic apparatus protein 1 | -4.06 | 0.00042 |
| *TMEM38A* | Trimeric intracellular cation channel type A | 4.15 | 0.00081 |
| *IARS1* | Isoleucine-tRNA ligase, cytoplasmic | 3.92 | 0.00098 |
| *RLN3* | Relaxin-3 | -4.24 | 0.0012 |
| *NPW* | Neuropeptide W | -3.32 | 0.0020 |
| *Rm15856d1426225t5* | Unknown | 3.59 | 0.0046 |
| *LOC120989780* | Cell wall protein IFF6-like | 5.77 | 0.0068 |
| *NUAK1* | NUAK family SNF1-like kinase 1 | -3.64 | 0.0106 |
| *Rm244708d4312373t2* | Unknown | -4.09 | 0.0106 |
| *KL* | Klotho | 3.88 | 0.0106 |
| *Rm56449d6040153t4* | Unknown | 5.18 | 0.0112 |
| *KIF3C* | Kinesin-like protein KIF3C | 3.46 | 0.0115 |
| ***EPB41L4A*** | Erythrocyte membrane protein band 4.1-like protein 4A | -3.52 | 0.0115 |
| *RCAN2* | Calcipressin-2 | -3.49 | 0.0115 |
| *TFR2* | Transferrin receptor protein 2 | 6.73 | 0.0115 |
| *TUBA* | Tubulin alpha chain | -4.89 | 0.0115 |
| *PRKAG2* | 5'-AMP-activated protein kinase subunit gamma-2 | 4.38 | 0.0141 |
| *GIGYF1* | PERQ amino acid-rich with GYF domain-containing protein 1 | -3.53 | 0.0144 |
| ***LRRK2*** | Leucine-rich repeat serine/threonine-protein kinase 2 | 5.02 | 0.0144 |
| *LRFN1L* | Leucine-rich repeat and fibronectin type III domain-containing protein 1-like protein | 3.40 | 0.0147 |
| *RAB5B* | Ras-related protein Rab-5B | -3.39 | 0.0466 |
| *MMP9* | Matrix metalloproteinase-9 | 5.02 | 0.0466 |

## Discussion

In this study, we set out to investigate transcriptomic changes in cane toads’ brains associated with expansion across its Australian invasive range. We found approximately five thousand DEGs between the Hawai’ian source population and the Australian invasive population. Therefore, extensive transcriptomic differences exist between the source and Australian population, despite a general similarity in genetic composition between source and range-core populations (F_ST_ = 0.04; Selechnik et al., 2019a). These differences in gene expression patterns likely reflect abiotic and biotic differences between the Hawai’ian source and Australian populations. Interestingly, more genes were upregulated than downregulated in the Australian invasive population than in the Hawai’ian source population. A general increase in brain gene expression in the Australian invasive population, particularly in genes involved in metabolism and mitochondrial function, might indicate a response to a novel and potentially more stressful environment (Kosmala et al., 2020; Tingley et al., 2014).

In contrast, the brain transcriptome of toads was remarkably similar across the entire Australian invasive range. Intermediate and range-front populations showed only a few dozen DEGs compared to the range-core population, and no significant difference in gene expression with each other. Looking at the overall transcriptomic response across all sites ordered along an east-to-west transect showed essentially the same result. While a few more DEGs were identified, the overall gene expression pattern remained consistent along the entire transect. This pattern confirms that the similarity in brain transcriptomes identified between range-core, intermediate and range-front populations is not an artefact of arbitrarily defined populations, but rather a biological feature of the Australian cane toad invasion. The strong geographic structuring of transcriptomic differences on either side of the Australian Great Dividing Range suggests that most of the divergence in Australian cane toads’ gene expression occurred early during the invasion process. The lack of gene expression differences observed between intermediate and frontal populations is compatible with their genetic structure, because these two populations are part of the same genetic cluster (Selechnik et al., 2019).

Likewise, the variability in brain gene expression was very limited across the Australian invasive range, with only 14 to 29 differentially dispersed genes between any two populations. Further, we failed to detect any sign of a change in gene expression variability from range-core to range-front populations, and therefore find no genome-wide support for changes to plasticity across this range. Although variability in gene expression can affect phenotypic differences (Ecker et al., 2018), this factor does not seem to have been important in cane toads.

The overall conservatism in brain gene expression seems to be at odds with the magnitude of phenotypic differences observed within the Australian invasive range (Brown et al., 2015; Brown et al., 2013; Gruber et al., 2017; Hudson et al., 2016; Hudson et al., 2018; Llewelyn et al., 2010; Phillips et al., 2006; Pizzatto et al., 2017; Tingley et al., 2012; Urban et al., 2008). Nonetheless, invasiveness may sometimes be promoted by a small number of genes (Bock et al., 2015), and minor changes in brain transcriptomes can underlie substantial differences in behaviour (Renn & Schumer, 2013; Saul et al., 2017). Moreover, other studies have reported low divergence in brain gene expression between populations with substantial morphological, behavioural and physiological differences, e.g. between domesticated and wild mammals (Albert et al., 2012), between domesticated and wild zebrafish (Drew et al., 2012), or between African and European fruit flies (Catalan et al., 2012). In other words, the extensive, and partially heritable, phenotypic differences seen between range-core, intermediate and range-front cane toad populations could be underlain by only a few key genes.

In favour of this hypothesis, some of the genes showing differential expression and/or dispersion between range-core, intermediate and range-front populations have putative functions related to behaviour. Lysosomal alpha-mannosidase (*MAN2B1*) is involved in neurocognitive functions such as learning and memory, and plays a role in motor function (D’Hooge et al., 2005; Damme et al., 2011). Calbindin (*CALB1*) is involved in locomotor behaviour (Barski et al., 2003). *CALB1* ko mice have lower anxiety-like behaviour, increased exploratory behaviour, and are less prone to exhibiting freezing behaviour (Harris et al., 2016). Leucine-rich repeat serine/threonine-protein kinase 2 (*LRRK2*) is involved in exploration behaviour and is linked with Parkinson’s Disease (Melrose et al., 2010). Mucolipin-3 (*MCOLN3*) is involved in locomotor behaviour, mutant mice showing erratic circling behaviour (Di Palma et al., 2002). Sodium-dependent neutral amino acid transporter (*SLC6A17*) mutations cause behavioural problems in humans (Iqbal et al., 2015). LIM homeobox transcription factor 1-alpha (*LMX1A*) is involved in memory and locomotor and olfactory behaviour, with mutations linked with Parkinson’s Disease (Laguna et al., 2015). Calcipressin-2 (*RCAN2*) is involved in locomotor behaviour, stress responses and memory (Miyakawa et al., 2003). Relaxin-3 (*RLN3*) is involved in exploratory behaviour, stress responses and the regulation of feeding behaviour (Smith et al., 2009). *MAN2B1*, *CALB1*, *LRRK2* and *MCOLN3* were all downregulated in intermediate populations (and range-front populations for *MAN2B1*) compared to range-core populations. Further, *MAN2B1*, *LRRK2* and *MCOLN3* were all under-dispersed in intermediate populations (and range-front populations for *MAN2B1*) compared to range-core populations. Both *LRRK2* and *MCOLN3* were under-dispersed in intermediate versus range-front populations. Both *SLC6A17* and *LMX1A* were upregulated in intermediate versus range-core populations. Both *RCAN2* and *RLN3* were under-dispersed in range-front versus intermediate populations. Therefore, the above genes may contribute to the behavioural shift in dispersal-related behaviour in toads from range-front and intermediate populations versus range-core populations.

A number of genes involved in the regulation of feeding behaviour also showed differential levels and/or variability of expression between range-core, intermediate and range-front populations. These include pro-opiomelanocortin (*POMC*) (Millington, 2007), nucleobindin-2 (*NUCB2*) (Dore et al., 2017), neuropeptide W (*NPW*) (Mondal et al., 2003) and agouti-related protein (*AGRP*) (Ollmann et al., 1997). *POMC* was downregulated in intermediate and range-front populations compared to range-core populations. *NUCB2* was upregulated in intermediate versus range-core populations, and was under-dispersed in range-front versus range-core populations. *NPW* was under-dispersed in range-front populations compared to range-core and intermediate populations. *AGRP* was over-dispersed in range-front versus intermediate populations. Toads from range-front areas have higher feeding rates, larger fat bodies, better body condition and faster growth (Brown et al., 2013). *POMC, NUCB2, NPW* and *AGRP* could thus also play a role in the phenotypic changes associated with range expansion in Australian cane toads.

Some genes showed a curvilinear pattern of gene expression along the range-core to range-front transect, with gene up- or downregulation in intermediate populations compared to both range-core and range-front populations. Curvilinear patterns across the invasive range have been found for cane toads’ spleen gene expression (Selechnik et al., 2019b), spleen mass, fat body mass, lungworm infection (Brown et al., 2015), and limb length (Hudson et al., 2016; Stuart et al., 2019). These curvilinear relationships with invasion history could be partly underlain by a ‘travelling wave’ density pattern, where higher population densities in intermediate populations compared to range-front and range-core populations cause a change in selection on dispersal-related traits (Brown et al., 2015). If so, this could explain some of the changes in brain gene expression that differentiate intermediate populations from range-front and range-core populations.

We must consider some potential limitations in our study that might explain our findings. Behaviour is a complex phenotype at the interplay between genetic and environmental factors. It is thought that specific gene regulatory networks interact with neuronal networks within particular brain regions to orchestrate behavioural responses to both internal (e.g., hormonal) and external (e.g., environmental) stimuli (Sinha et al., 2020). The low magnitude of gene expression changes that we observed in the Australian invasive population could thus be partially obscured by concomitant and opposing changes in gene expression across various brain regions (Nadler et al., 2006). Furthermore, heterogeneity in physiological and/or environmental conditions between individuals can confound any transcriptomic analysis, especially for individuals sampled in the wild (as in the present study). It is also possible that by sampling adult toads, we missed the critical developmental window during which key genes underlying behavioural plasticity are differentially expressed (Aubin-Horth & Renn, 2009). Finally, additional molecular mechanisms inaccessible with RNA-seq (such as post-translational modifications) might play an important role in behavioural changes (Cash et al., 2005).

Genetic differentiation is low across the cane toads’ Australian invasive range (Leblois et al., 2000; Lillie et al., 2014; Selechnik et al., 2019a; Slade & Moritz, 1998), but toads from range-front and intermediate populations form a distinct genetic cluster compared to range-core toads (Selechnik et al., 2019a). We found here that these genetic differences are mirrored by changes in gene expression on either side of the Australian Great Dividing Range. This phenomenon might be an example of rapid evolution, or a case of environmentally-induced variation (phenotypic plasticity). We note, however, that phenotypic plasticity can also lead to adaptive evolution (Ghalambor et al., 2007), e.g., through genetic assimilation (Pigliucci & Murren, 2003; West-Eberhard, 2003). If evolution is at play, does it act through shifts in selective regimes (adaptive evolution), or through non-adaptive processes (e.g., drift, spatial sorting, admixture), or through a combination of factors?

In conclusion, the modest differences that we have documented in brain gene expression along the invasive range of cane toads within Australia might trigger significant changes in the toads’ phenotypic traits, in particular in relation to dispersal behaviour. Common-garden experiments monitoring dispersal-related behaviour together with gene expression in the identified genes would provide valuable insight into the heritability, and thus evolvability, of these phenotypic changes.

## Acknowledgements

We thank Simon Ducatez, Cameron Hudson, Chris Jolly and Serena Lam for assistance in the field and Kate Buchanan for her invaluable support and contribution facilitating this project. This work was supported by the Australian Research Council (FL120100074 to RS, DE150101393 to LAR), Deakin University (CRGS #27701 to LAR) and the UNSW Scientia Program (to LAR).

## Data Accessibility

We have deposited the raw RNA-seq data to the National Center for Biotechnology Information (NCBI) Sequence Read Archive (BioProject PRJNA479937).

## Author Contributions

MFR and LAR conducted fieldwork. AJW, MFR and DS conducted lab work. BY, AJW and MFR conducted analyses. BY, AJW, MFR, DS, RS and LAR contributed to interpretation and writing.

---

**Figure S1.** Heatmap of correlation of gene expression for all pairwise combinations of samples. Colour depicts Pearson’s *r* correlation coefficient. Samples deemed as outliers are indicated with asterisks.

