## Supplementary material for "Brain transcriptome analysis reveals gene expression differences associated with dispersal behaviour between range-front and range-core populations of invasive cane toads in Australia": Table S1

**Table S1.** Location of samples. HI, Hawai’i; QLD, Queensland; NT, Northern Territory; WA, Western Australia.

| Sample | Coordinates | Location | State | Ecotype |
| --- | --- | --- | --- | --- |
| RMH051B | 19°57'N, 154°96'W | Paradise Park | HI | Source |
| RMH052B | 19°57'N, 154°96'W | Paradise Park | HI | Source |
| RMH059B | 19°57'N, 154°96'W | Paradise Park | HI | Source |
| RMH061B | 19°57'N, 154°96'W | Paradise Park | HI | Source |
| RMH075B | 19°94'N, 155°88'W | Mauna Lani Point | HI | Source |
| RMH077B | 19°94'N, 155°88'W | Mauna Lani Point | HI | Source |
| RMH079B | 19°94'N, 155°88'W | Mauna Lani Point | HI | Source |
| RMH081B | 19°94'N, 155°88'W | Mauna Lani Point | HI | Source |
| H3 | 21°42'N, 157°82'W | Haiku Gardens | HI | Source |
| H4 | 21°42'N, 157°82'W | Haiku Gardens | HI | Source |
| RMH008B | 21°42'N, 157°82'W | Haiku Gardens | HI | Source |
| RMH043B | 21°42'N, 157°82'W | Haiku Gardens | HI | Source |
| RMH047B | 21°42'N, 157°82'W | Haiku Gardens | HI | Source |
| H1 | 21°34'N, 158°08'W | Kapolei Regional Park | HI | Source |
| H2 | 21°34'N, 158°08'W | Kapolei Regional Park | HI | Source |
| RMH026B | 21°34'N, 158°08'W | Kapolei Regional Park | HI | Source |
| RMH027B | 21°34'N, 158°08'W | Kapolei Regional Park | HI | Source |
| RMH028B | 21°34'N, 158°08'W | Kapolei Regional Park | HI | Source |
| B31 | 17°08'S, 145°80'E | Gordonvale | QLD | Core |
| B32 | 17°08'S, 145°80'E | Gordonvale | QLD | Core |
| B33 | 17°08'S, 145°80'E | Gordonvale | QLD | Core |
| B34 | 17°08'S, 145°80'E | Gordonvale | QLD | Core |
| B29 | 16°25'S, 145°32'E | Daintree | QLD | Core |
| B30 | 16°25'S, 145°32'E | Daintree | QLD | Core |
| B35 | 16°25'S, 145°32'E | Daintree | QLD | Core |
| B36 | 16°25'S, 145°32'E | Daintree | QLD | Core |
| B25 | 18°21'S, 142°24'E | Croydon | QLD | Intermediate |
| B26 | 18°21'S, 142°24'E | Croydon | QLD | Intermediate |
| B27 | 18°21'S, 142°24'E | Croydon | QLD | Intermediate |
| B28 | 18°21'S, 142°24'E | Croydon | QLD | Intermediate |
| B21 | 17°85'S, 139°63'E | Burketown | QLD | Intermediate |
| B22 | 17°85'S, 139°63'E | Burketown | QLD | Intermediate |
| B23 | 17°85'S, 139°63'E | Burketown | QLD | Intermediate |
| B24 | 17°85'S, 139°63'E | Burketown | QLD | Intermediate |
| B5 | 16°67'S, 135°80'E | Cape Crawford | NT | Intermediate |
| B6 | 16°67'S, 135°80'E | Cape Crawford | NT | Intermediate |
| B7 | 16°67'S, 135°80'E | Cape Crawford | NT | Intermediate |
| B8 | 16°67'S, 135°80'E | Cape Crawford | NT | Intermediate |
| B1 | 14°92'S, 133°07'E | Mataranka | NT | Intermediate |
| B2 | 14°92'S, 133°07'E | Mataranka | NT | Intermediate |
| B3 | 14°92'S, 133°07'E | Mataranka | NT | Intermediate |
| B4 | 14°92'S, 133°07'E | Mataranka | NT | Intermediate |
| B9 | 15°64'S, 130°47'E | Timber Creek | NT | Intermediate |
| B10 | 15°64'S, 130°47'E | Timber Creek | NT | Intermediate |
| B11 | 15°64'S, 130°47'E | Timber Creek | NT | Intermediate |
| B12 | 15°64'S, 130°47'E | Timber Creek | NT | Intermediate |
| B17 | 15°91'S, 128°18'E | Durack River | WA | Front |
| B18 | 15°91'S, 128°18'E | Durack River | WA | Front |
| B19 | 15°91'S, 128°18'E | Durack River | WA | Front |
| B20 | 15°91'S, 128°18'E | Durack River | WA | Front |
| B13 | 18°22'S, 127°67'E | Halls Creek | WA | Front |
| B14 | 18°22'S, 127°67'E | Halls Creek | WA | Front |
| B15 | 18°22'S, 127°67'E | Halls Creek | WA | Front |
| B16 | 18°22'S, 127°67'E | Halls Creek | WA | Front |
