## Supplementary figures and images for "Brain transcriptome analysis reveals gene expression differences associated with dispersal behaviour between range-front and range-core populations of invasive cane toads in Australia"

### Figure S1

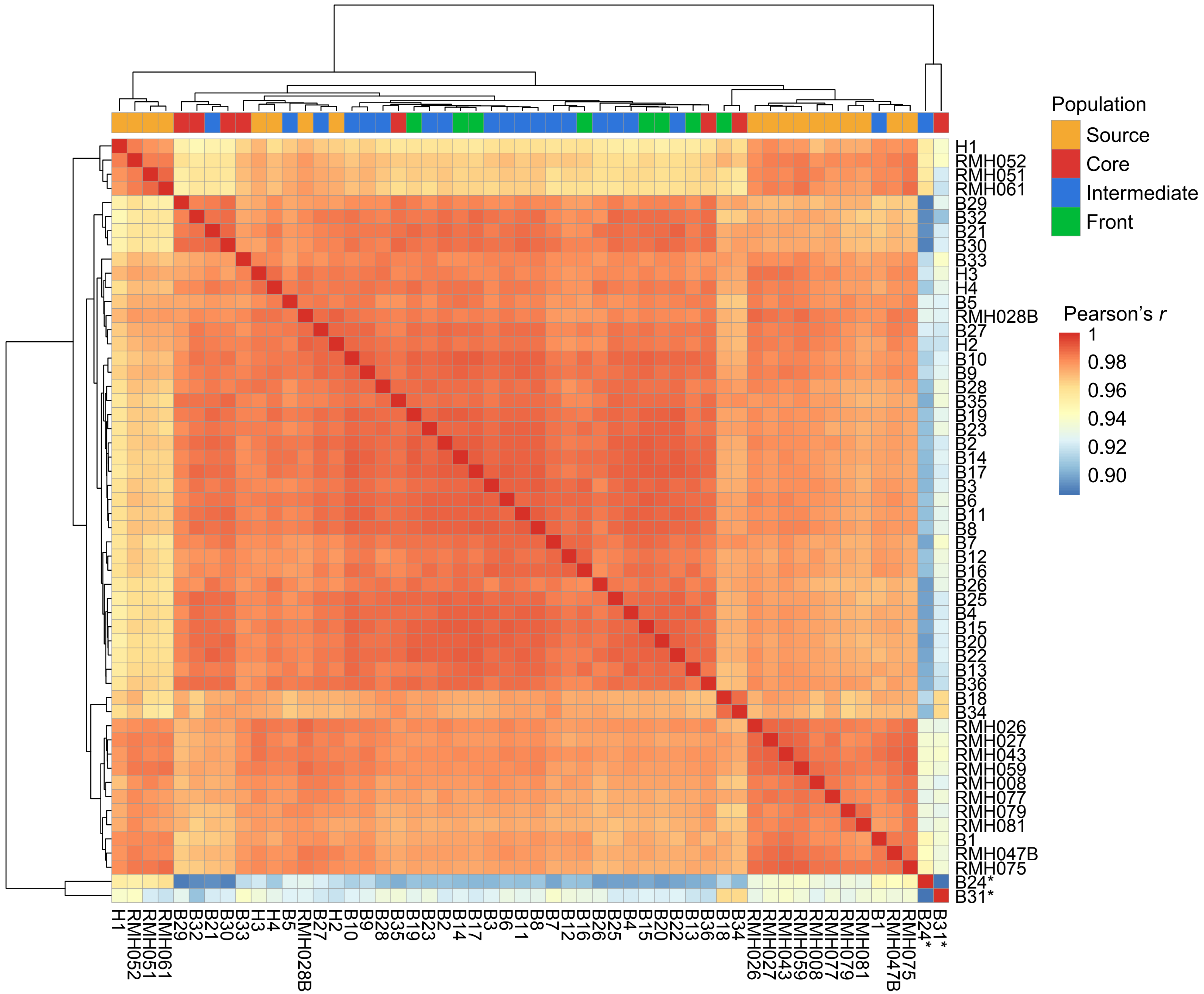
